# SOYBEAN RESPONSE TO INITIAL STAGE OF *Fusarium virguliforme* INFECTION

**DOI:** 10.1101/2022.03.07.483270

**Authors:** María Lorena Giachero, Nathalie Marquez, Leandro Ismael Ortega, Daniel Adrian Ducasse

## Abstract

*Fusarium virguliforme* causes the Sudden Death Syndrome, an important disease in soybean crops. In this work, we investigated the defensive response mechanisms in soybean root, at cell wall level, during *F. virguliforme* infection using an in vitro culture system. We measured total root lignin content by the acetyl bromide method and estimated the in-situ lignin and suberin deposition by confocal microscopy on local and systemic root tissues, i.e. adjacent and distant to the pathogen entry site respectively. Moreover, the expression dynamics of phenylalanine ammonia lyase (PAL), shikimate/quinate hydroxycinnamoyltransferase (HCT) and cinnamyl alcohol dehydrogenase (CAD) was evaluated by real-time quantitative PCR. The results showed that, although the most significant increment of lignin deposition was observed in the epidermal cells of local tissues, *F. virguliforme* also induced lignin deposition changes in a sistemic fashion. In fact, inoculated plants presented a higher deposition of lignin in hypodermis and cortex than the control ones, independently of the distance from the inoculum source, while suberin deposition was higher in local zones. Moreover, the gene expression analysis showed an up-regulation of PAL, HCT and CAD genes after the inoculation with the pathogen, which correlates with the cell wall modifications observed in the local tissues. The results presented here suggest that the increase in lignin and suberin deposition during soybean root/*F. virguliforme* interaction is probably a strategy not only to stop the pathogen entrance, but to provide the plant more time to prepare its defences as well.

## 1. INTRODUCTION

In 2020, the world soybean production reached around 350 million tons over 126.9 million ha. Climate, land extension, and soil conditions turn Brazil (121.8 million tn), United States (112.5 million tn) and Argentina (48.8 million tn) the major world soybean producers (FAOstat 2021). On the other hand, these geographical regions may have optimal conditions for the occurrence of soybean diseases and consequently yield losses (Wrather et al. 2010). Sudden Death Syndrome (SDS) is one of these diseases and it is ranked as the most destructive for this crop worldwide (Wrather et al. 2010). SDS is caused by a complex of at least four species of *Fusarium* sp.: *F. virguliforme, F. tucumaniae, F. brasiliense* and *F. crassistipitatum* (Aoki et al. 2005, 2012). However, *F. virguliforme* is the most geographically distributed (Spampinato et al. 2021). The colonization of these fungi is generally restricted to roots, however SDS causes both root and leaf symptoms. Root symptoms include crown rotting and vascular tissue discoloration, while leaf symptoms consist in typical chlorotic interveinal spots that can coalesce and eventually become necrotic (Roy 1997; Rupe et al. 2001). The toxins produced by the pathogen in roots are responsible for foliar symptoms (Brar et al. 2011; Abeysekara and Bhattacharyya 2014). The severity of foliar symptoms of SDS is associated with soil temperature (22 to 24°C) and soybean age, given plants are more susceptible to infection at early stages of development (Gongora-Canul et al. 2011).

*F. virguliforme* is a hemi-biotrophic pathogen (Roy et al. 1997), initially behaving as a biotrophic and later switching to a necrotrophic phase (Iqbal et al. 2005). Navi et al. (2008) showed that *F. virguliforme* forms appressoria and infection pegs to penetrate the host root. In a comparative transcriptome analysis of *F. virguliforme* infection process, Sahu et al. (2017) found an upregulation of genes related to the degradation of antimicrobial compounds such as phytoalexin and glycoline during the early stages of root tissue infection. Meanwhile, in late infection stages, many genes encoding hydrolytic and catalytic enzymes were up-regulated (Sahu et al. 2017). An important component of pathogen attack are the cell wall degrading enzymes (CWDEs). A recent study has shown that also it is likely that *F. virguliforme* employs CWDE as pathogenicity factors (Islam et al., 2017). Plants can prevent pathogen infection by inactivating CWDEs, through secreting inhibitor proteins (PIPs) (Chang et al. 2016; Misas-Villamil and Van der Hoorn 2008). Interestingly, two xylanases of *F. virguliforme* related to cell wall degradation, were detected as potential targets for exogenous PIPs (Chang et al. 2016).

Several studies have been carried out to characterize the plant response to *F. virguliforme* colonization. During soybean infection, genes related to defense, signal recognition and transduction, cell wall synthesis and metabolic processes were differentially expressed (Iqbal et al., 2005; Nagki et al., 2016; Marquez et al., 2019; Radwan et al., 2011). A large number of these up-regulated genes encode cell wall and plasma membrane related proteins (Ngaki et al., 2016). Curiously, even though phenylpropanoid biosynthesis was the metabolic pathway with the highest number of differentially expressed genes, Marquez et al. (2019) observed the downregulation of a considerable number of cell wall and signalling-related genes in presence of *F. virguliforme*. Similarly, previous soybean transcriptome studies showed an induction of the phenylpropanoid pathway after inoculation with *Fusarium solani* f. sp. *Glycines* (Lozovaya et al., 2004; Iqbal et al., 2005). Apparently, the soybean roots of partially resistant genotypes display high rate of lignification immediately post infection (Lozovaya et al. 2006). This result shows that lignin is playing a key role in the defence against this pathogen. Moreover, Iqbal et al. (2005) suggest that, in addition to the phenylpropanoid pathway, the partially resistant soybean genotypes to SDS are capable of recognizing the pathogen through receptors similar to kinases (RLK) as an early defence mechanism.

The structure and composition of plant cell wall is dynamic, and can be remodeled during development or in response to the environment (Liu et al. 2018). The cell wall, in addition to being a physical barrier against pathogen invasion, has been proposed as a monitor system. Plants can detect modifications in the cell wall and trigger immune responses, through protein kinases and calcium signaling (Wolf 2017). Lignin is one of the main and most widespread cell wall strengtheners in the Tracheophyta (Zhong et al. 2011). Lignin biosynthesis is often induced at the site of pathogen attack providing a physical barrier against infection (Bonello et al. 2003; Buendgen et al. 1990). Likewise, suberin also constitutes a pathogen-induced defence response. Suberization occurs in specific sites of the roots (Thomas et al. 2007) forming a physical barrier and preventing water loss through exposed and injured tissues (Kolattukudy 2001; Franke and Schreiber 2007).

The main purpose of this work was to investigate, at cell wall level, the defensive response mechanisms in soybean root during the early steps of *F. virguliforme* infection using an in vitro culture system. Therefore, the present study aims were 1) to quantify total root lignin content, 2) to observe the pattern of in-situ lignin and suberin deposition and distribution in two roots sampling zones: local (adjacent to the pathogen entry - FvA) and systemic (distant to the pathogen entry - FvD) and 3) to evaluate the expression levels of lignin-related biosynthetic pathway genes (local and systemic).

## 2. MATERIALS AND METHODS

### 2.1 Biological Material

*F. virguliforme* (Syn. *F. solani* f. sp. *glycines*) (*Fusarium virguliforme* O’Donnell & T. Aoki) obtained from soybean (*Glycine max*) in Argentina (Buenos Aires, San Pedro) (Aoki et al. 2005) was grown on Potato Dextrose Agar (PDA) (Scharlau Chemie S.A, Barcelona, Spain) at 25°C in the dark for seven days.

Soybean seeds [*Glycine max* (L.) Merr.] cv SPS 4×4RR (Syngenta) were surface-disinfected following protocol descript by Giachero et al. (2017). After disinfection, groups of 6 seeds were germinated per Petri dish (Ø145 mm) with 1% (w/v) water-agar supplemented with 0.8% sucrose. The Petri dishes were kept in the dark at 25°C for 4 days, and then exposed to light for 24 h (average photosynthetic photon flux of 225 μmol m−2 s−1).

### 2.2 Experimental design

Soybean plantlets (5 days old) were transferred to an *in vitro* system containing Modified Strullu-Romand (MSR) medium (Declerck et al. 1998) without sucrose and vitamins, and solidified with 3 g l^-1^ Phytagel (Sigma-Aldrich, St. Louis, USA). Plantlets roots were placed on the surface of the medium and the shoots were let extrude the Petri dish through a hole in the lid (Giachero et al. 2017). These *in vitro* systems were incubated in a growth chamber at 25ºC, 70% of relative humidity, and 16 h/day photoperiod with a 300 μmolm^−2^s^−1^ photosynthetic photon flux. The dishes were covered to avoid light incidence on the roots. After fifteen days, the *in vitro* systems were randomly divided into two groups corresponding with two treatments: one control without *F. virguliforme* inoculation (C) and the other one, inoculated with *F. virguliforme* (Fv). The inoculation protocol was performed as described by Giachero et al (2017) using a 7 days old plug from a *F. virguliforme* culture in PDA. Root samples were taken from both treatments at 24, 48, 72 and 96 hours post-inoculation (hpi) for the determinations described in the next sections.

### 2.3 Root lignin content determination by Acetyl bromide method

This method allows an in-bulk determination of the lignin content of whole plant organs. To determine lignin content in soybean roots, the acetyl bromide protocol descript by Moreira-Vilar et al. (2014) was used with modifications.

The absorbance of the samples was measured at 280 nm. A standard curve was generated using Kraft lignin (Sigma-Aldrich®) and the results were expressed as lignin (mg) per cell wall (g). Four biological replicates were considered for each treatment at every time point.

### 2.4 *In-situ* lignin and suberin estimation by confocal microscopy

*In-situ* lignin and suberin deposition was estimated as described by Hutzler et al. (1998) taking advantage of the auto-fluorescent properties of these biopolymers (Vaughn and Lulai 1991).

Root segments of 2 cm in length, were sampled from a site near to the pathogen entry (FvA, local) and distant from that point (FvD, systemic) (Figure 1a). The segments corresponded to the root maturation zone (Figure 1b), i.e. 1 cm from the root apex. Transverse free-hand sections were performed at 5mm intervals along the fragment, which were immediately fixed in FAA (formaldehyde, acetic acid, EtOH, 2:1:17) for half an hour and mounted onto glass-slides with a drop of 50% glycerol. Observations were done in a Nikon Eclipse CS1 spectral confocal microscope, using a 400 nm diode laser at 10% power as light source. In preliminary experiments, the emission was analysed with the spectral detector (28 channels, 10 nm resolution) in the different zones to be examined: epidermis (e), hypodermis (h), cortex (c) and endodermis (en, Figure 1c). Differential spectra attributed to lignin and suberin were successfully discriminated (data not shown). From these results, the correspondence between emission peaks of each biopolymer and the emission filters range of the conventional confocal mode detectors was observed. Thus, this acquisition mode was used henceforth for the rest of the analyses. Auto-fluorescence emission, due to lignin and suberin, was imaged through BP450/35 (“blue” channel) and BP605/75 (“red” channel) emission filters, respectively. Images were acquired setting 10 μs of laser dwell-time and a 1024×1024 dpi resolution. Average pixel gray value in each channel was measured inside delimited areas, using the open source software FIJI (Schindelin et al. 2012). Four biological replicates per treatment in each time point of sampling, were used in each case. This experiment was performed twice.

**Figure 1.**
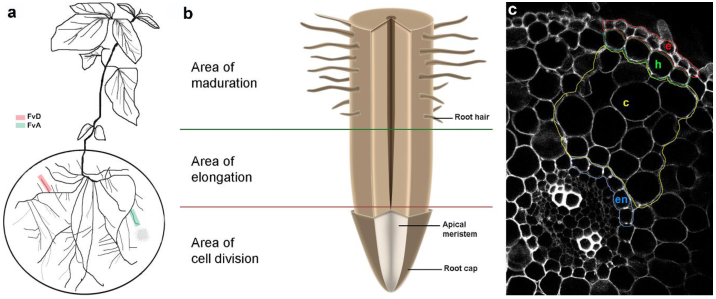
(**a**) Schematic representation of the in vitro culture system used to grow soybean plants in Petri dishes with Phytagel medium. The figure shows the two sampling sites of the root system, adjacent (FvA) and distant (FvD) to the pathogen inoculation point. (**b**) Schematic view (not at scale) of the different zones of the root tip. Segments from the maturation zone were sampled for sectioning and observation. (**c**) Representative image showing the four evaluated areas: epidermis (e), hypodermis (h), cortex (c) and endodermis (en).

### 2.5 Gene expression analysis by real-time quantitative RT-PCR

Root segments of 2 cm in length were sampled from C (control), FvA (local) and FvD (systemic) at 24, 48, and 72 hpi with *F. virguliforme*.

Total RNA was extracted using the Trizol® reagent (Invitrogen, Carsbad, USA) according to manufacturer’s instructions with an additional chloroform purification step. All samples were treated with DNAse I Amplification Grade (Invitrogen Life Technologies) according to the manufacturer’s instructions. Concentration and purity of total RNA were determined (A260/280 absorbance ratio) using a NanoDrop®-ND 1000 UV-Vis Spectrophotometer (NanoDrop Technologies, Wilmington, USA).

Real-time quantitative RT-PCR (qPCR) was performed using the Rotor Gene 6000 with the iTaqTM Universal SYBR® Green Supermix (BIORAD, catalog 172-5151) and specific primers for the following genes: Phenylalanine ammonia lyase 1 (PAL 1), Cinnamyl alcohol dehydrogenase (CAD), Shikimate O-hydroxycinnamoyltransferase (HCT) (Supplementary table 1). The thermal cycling protocol was: 50ºC for 10 min, 95ºC for 1 min, amplification and quantification cycles repeated 40 times 95ºC for 15 s, 61ºC for 30 s, followed by melting curve data collection to check for non-specific amplification and primer dimers. Three independent biological replicates, each consisting of a pool of two soybean root samples, with two technical replicates were analyzed per treatment. Normalization was achieved using the SKIP16 reference gene (Hu et al. 2009), which showed to be stable under the tested conditions. Relative gene expression was calculated using the 2^−ΔΔCt^ method (Livak and Schmittgen, 2001). The amount of target transcripts in control roots (C) was considered as basal level during expression analysis.

### 2.6 Assessment of colonization by *F. virguliforme*

The root samples for assessment of infection by *F. virguliforme* were taken at the end of assay (i.e. 96 hpi). Colonization by *F. virguliforme* was followed under the microscope (Nikon labophot-2, Japan) at 10–40× magnification. Roots were cleared in 10% KOH at room temperature for 3 h, rinsed with distilled water, bleached and acidified with HCl 1% and stained with Trypan blue 0.2% at room temperature overnight.

### 2.7 Statistical analysis

Data were subjected to analysis of variance (ANOVA) (p≤ 0,05 level of significance). The LSD Fisher post-hoc test was used to discriminate differences in lignin and suberin content results (spectrophotometry and microscopy) and the DGC test for the qPCR experiments. An *α*=0.05 was used in all cases. All the analyses were performed using the INFOSTAT software (Di Rienzo et al. 2008).

## 3. RESULTS

### 3.1 Lignin and suberin content

Lignin content in the total root system, determined by the acetyl bromide method, did not show significant differences between treatments during the evaluated time points (Figure 2). Since this approach could overlook differences in lignin deposition at cell wall level, the root maturation zone was examined by cross-sections (Figure 3). This analysis showed that the amount of lignin was different between treatments. Using a semi-quantitative estimation by Average Grey Value (AGV) of microscopy images, we observed that *F. virguliforme* inoculation induced changes in lignin deposition. Both FvA and FvD zones showed a significant increase in lignin content with respect to non-inoculated root (C) at 48 and 96 hpi with an apparent higher increase in FvD at 48 hpi (Figure 3a,c). Irrespective to time, the root epidermis (e) showed the highest AGV in all treatments with a significant increase in the adjacent zone of the inoculated plants (Figure 3b). Both hypodermis (h) and cortex (c) of inoculated plants, from either FvA and FvD segments, showed higher AGVs with respect to controls (Figure 3b). No differences were detected in the endodermis (data not shown). Suberin was detected both in epidermis and hypodermis cells exhibiting auto-fluorescence in tangential and radial cell walls. This auto-fluorescence was higher in epidermal than hypodermal cells (Figure 4b). AGVs were significantly higher in FvA zones only at 24 hpi (Figure 4a,c). The figure 4c shows a detailed image of both epidermis and hypodermis, where a high difference in the auto-fluorescence emission due to suberin (“red” channel) in FvA zones 24 hpi, was observed. As with lignin, no differences were detected in the endodermis (data not shown).

**Figure 2.**
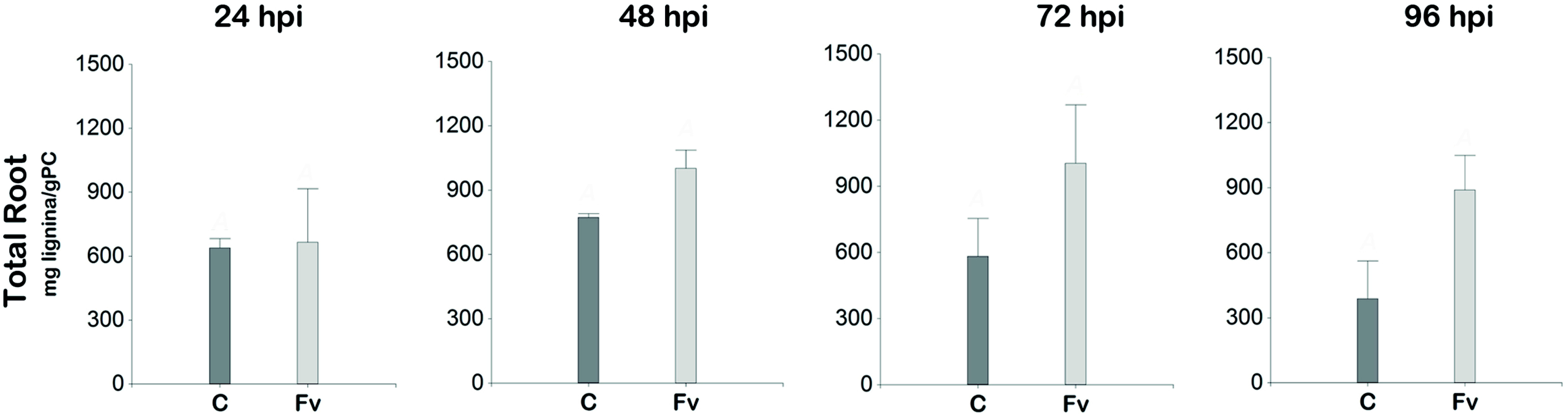
Lignin content of soybean root determined by the acetyl bromide method. C: control, Fv: roots of soybean inoculated with *F. virguliforme*. The results were expressed as mg lignin g-1 cell wall. Mean values ± SE (n = 3) marked with different letters are significantly different (p≤ 0,05 ANOVA and LSD Fisher *α*=0,05)

**Figure 3.**
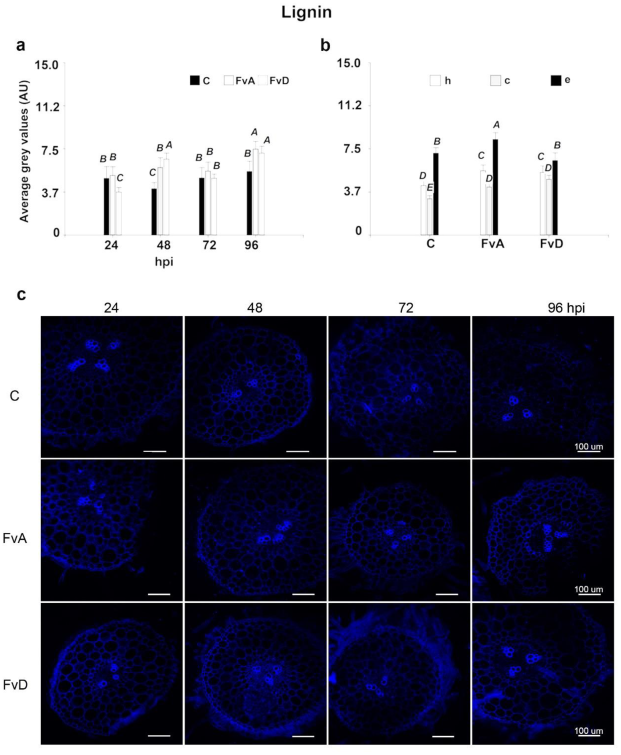
Lignin deposition in soybean roots (**a** and **b**) expressed like average grey values (AU: Arbitrary Units). FvA: adjacent to the Fv inoculation; FvD: distant area to the pathogen inoculation point; C: control; hpi: hours post inoculation. Evaluated areas: epidermis (e), hypodermis (h), cortex (c). Mean values ± SE (n = 3) marked with different letters are significantly different (p≤ 0,05 ANOVA and LSD Fisher α=0,05). Imaging of transverse free-hand sections in soybean roots (**c**). Observations were done in a Nikon Eclipse CS1 spectral confocal microscope. Auto-fluorescence emission due to lignin was imaged through BP450/35 (“blue” channel) emission filters.

**Figure 4.**
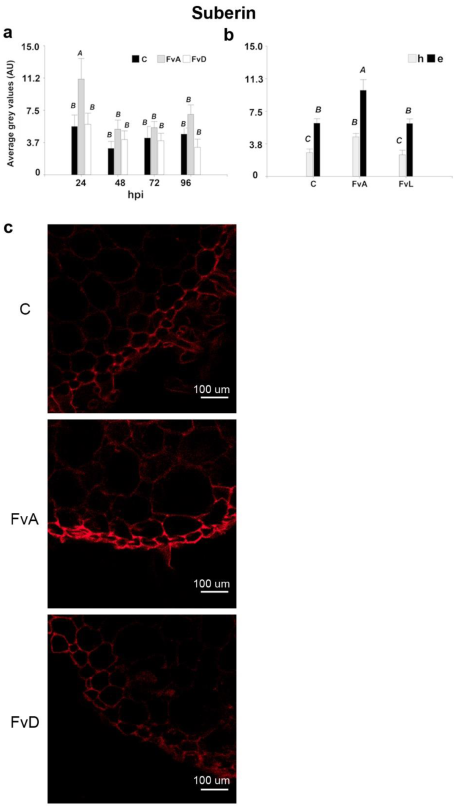
Suberin deposition in soybean roots (**a** and **b**) expressed like average grey values (AU: Arbitrary Units). FvA: adjacent to the Fv inoculation; FvD: distant area to the pathogen inoculation point; C: control; hpi: hours post inoculation. Evaluated areas: epidermis (e), hypodermis (h), cortex (c). Mean values ± SE (n = 3) marked with different letters are significantly different (p≤ 0,05 ANOVA and LSD Fisher α=0,05). In b, interaction between treatments and sampling zones was not significant (p ≤ 0,05). Nevertheless, comparisons of all the means involved in the interaction were assessed. Detailed image of epidermis and hypodermis in transverse free-hand sections in soybean roots (**c**) shows a difference in Auto-fluorescence emission due to suberin at 24 hpi. The image was imaged through BP605/75 (“red” channel) emission filters.

### 3.2 Gene expression analysis

The expression dynamics of three genes (PAL, HCT and CAD) was evaluated by real-time quantitative PCR in both treatments: inoculated with *F. virguliforme* (Fv) and non-inoculated (C) soybean roots at 24, 48, and 72 hpi. Two sampling areas were evaluated in the root system: adjacent (FvA) and distant (FvD) to the pathogen inoculation point.

Interaction between treatments, time (hpi) and sampling zones, was not significant (p ≤ 0,05) for the analysed genes. Nevertheless, comparison of all the means involved in the interaction was assessed.

The results (Figure 5) showed that PAL 1, HCT and CAD genes were significantly induced by *F. virguliforme*. However, the expression levels were similar among the three post-inoculation times analysed. Moreover, when soybean root sampling areas were compared, results showed that transcriptional changes of PAL 1, HCT and CAD genes were significantly higher in the adjacent area (FvA) than the distant area (FvD) to the pathogen inoculation point.

**Figure 5.**
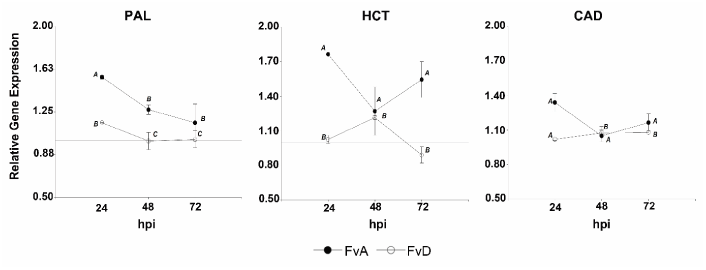
Quantitative RT-PCR of genes involved in the lignin biosynthesis pathway (PAL, HCT and CAD) in 2-week-old soybean roots in response to *Fusarium virguliforme* inoculation. Two sampling areas were evaluated in the root system: adjacent (FvA: dark spot) and distant (FvD: clear spot) to the pathogen inoculation point at 24, 48, and 72 hpi with *F. virguliforme*. Data are means of three biological replicates (n=3). The mean values and standard errors were calculated from two technical replicates. Transcript levels are expressed relative to non-inoculated soybean plantlets (C). Different letters indicate significant differences (DGC Alfa=0,05)

### 3.3 *F. virguliforme* colonization

The hyphae of *F. virguliforme* reached the soybean root surface at 48 h after inoculation. Following this first contact, hyphae development within the root could not be confirmed by microscopic observation and soybean roots did not show any disease symptoms.

## 4. DISCUSSION

In the current study, using an in vitro culture system, cell wall modification in soybean root composition was analysed during the early steps of *F. virguliforme* infection. Previous studies reported that different types of biotic stresses can raise lignin and suberin deposition in cell walls to block pathogen invasion and to reduce the host susceptibility (Moura et al. 2010; Thomas et al. 2007). Therefore, in a first assay, the amount of lignin was evaluated in the total root. Results showed a trend of increased lignin deposition in inoculated plants by the time course. However, this correlation could not be confirmed statistically when the roots were evaluated as a whole organ. Lignin deposition is a tissue specific process, occurring in key plant development processes as the conductive tissue maturation, which is fundamental for the proper function of xylem vessels (Barros et al. 2015). Moreover, several reports, have suggested that lignin biosynthesis is induced at the site of pathogen infection providing a physical barrier (Bonello et al. 2003; Buendgen et al. 1990) which may limit further growth and / or confine invading pathogens (Menden et al. 2007; Smit and Dubery, 1997; Wuyts et al. 2006). Thus, it is possible that this localized lignin deposition, induced by the defence response, be diluted in the whole organ analysis. This led us to evaluate the lignin deposition in situ, at the maturation zone, where cells no longer elongate (Figure 1b). It is in this root segment where lignin deposition is triggered as a defence reaction when a root pathogen approximates or pre-penetrates a root (Mandal and Mitra, 2007). In this study, taking advantage of the auto-fluorescent properties of cell wall biopolymers, confocal microscopy was used to estimate lignin and suberin content after *F. virguliforme* inoculation in local and systemic root tissues. Lignin deposition occurred mainly at the epidermis in the local tissues (FvA). In addition, hypodermis and cortex showed a higher lignin deposition in the inoculated roots compared to the non-inoculated ones, regardless of the distance to the inoculation point (FvA and FvD; Figure 3b). Interestingly, plants deposited lignin systemically in response to the fungus. Histochemical studies in *Brassica napus* revealed that the phenolics compounds and lignin deposition seems to be involved in vascular obstruction and reinforcement of tracheary elements during the early defence against Verticillium longisporum (Eynck et al. 2009). Particularly in the case of SDS, cell wall strengthness by lignin deposition is very important since *F. virguliforme* hydrolytic enzymes may play a major role in root necrosis during late infection stages (Sahu et al. 2017). A significant lignin deposition increase was detected in inoculated plants (FvA and FvD) at 48 and 96 hpi (Figure 3a). Curiously, although a gradual lignin deposition increase over time was expected, results showed a return to basal levels (not significant differences) at 72 hpi. This oscillation may be a consequence of the high variability in maturation stages of the experimental units (the segments of the maturation zone) which must be destructively sampled for the microscopic observation. An in-vivo following up of lignin deposition along time in each individual root, would have avoided this artifact.

In this work, suberin was detected in every evaluated condition. Suberin is constitutively deposited in root endodermal and peridermal cell walls (Vishwanath et. al, 2015). However, a high suberization level in the local tissues (FvA), in both the epidermis and the hypodermis cells, was evidenced. Confocal microscopy results showed that after fungus inoculation, suberin was first deposited, followed by lignin. Interestingly, this increment in suberitation was detected before the pathogen reached the root surface (i.e. 24 hpi), while lignification started 48 hours later, when *F. virguliforme* contacted the root surface. Genome sequencing of *F. virguliforme* identified putative pathogenicity genes. Among them there were genes that may be associated with cell wall degradation in root tissues (Srivastava et al 2014). Results presented in this study, suggest that the plant is capable of detecting the pathogen secreted molecules during the early infection stage and inducing suberin deposition near the *F. virguliforme* entrance to delay fungus infection in the roots. There are several reports of pathogen eliciting suberin deposition in the host cell walls within and around the infection site, limiting the spread (Kolattukudy and Espelie, 1989). Thomas et al. (2007) demonstrated that preformed suberin reduced soybean susceptibility to Phytophthora sojae, suggesting a relationship between suberin deposition and disease resistance. The suberization might be an interesting target to improve plant resistance against pathogens. In addition, suberin associated compounds (i.e. wax or phenolics components) may have antifungal properties (Lulai and Corsini 1998).

A considerable number of genes encoding cell wall and plasma membrane proteins were induced in soybean roots after *F. virguliforme* infection (Ngaki et al. 2016). Particularly, the phenylpropanoid pathway genes are highly expressed in biotic stress conditions (Yadav et. at. 2020). In this work we evaluated the expression dynamics of three of those genes: PAL, HCT and CAD. As result, the expression induction by *F. virguliforme* of the three evaluated genes was observed even before the root-hyphae contact (24 hpi) and correlated with the cell wall modifications in the adjacent area (FvA). Marquez et al. (2019) demonstrated that the phenylpropanoid biosynthesis pathway showed the highest number of differentially expressed genes upon *F. virguliforme* infection. Phenylpropanoid metabolism pathway generates lignin precursors, flavonoids and other aromatic metabolites like coumarins, volatile phenolic compounds, or tannins (Cheynier et al. 2013). This suggests that the premature activation of the phenylpropanoid biosynthesis pathway may result in the plant cell resistance to the pathogen entry through incrementing a wide variety of precursors, as structural and/or signalling molecules. A similar response was observed in resistant ginger against Pythium, a necrotrophic soil-borne that causes soft-rot disease (Geetha et al. 2019). In conclusion, results presented in this work provide further evidence of an induced increment of lignin and suberin deposition specifically in the infection site of *F. virguliforme* in soybean roots. This event is probably part of the plant defence response intended to avoid the pathogen income or at least to delay pathogen invasion, giving the host more time to prepare the defence strategy. Strikingly, these results suggest that the roots not only trigger cell wall strengthening near the pathogen entry, but also in distant areas where strengthening of hypodermis and cortex was observed.

## ACKNOWLEDGEMENTS

This study was supported by the INTA’s research funds (PNPV-1135024). The authors would like to thank Dr. Gaston Vaghi Medina for his helpful suggestions and critical reading of the manuscript.

## AUTHOR CONTRIBUTIONS

MLG, NM, LO and DAD conceived and planned the experiments. MLG carried out the experiment, took and prepared the samples for confocal microscope observation, analysed the data and took the lead in writing the manuscript. NM and LO contributed to the analysis of the results and to the writing of the manuscript. LO contributed to sample preparation and confocal microscopy observations.

All authors provided critical feedback and helped shape the research, analysis and manuscript.

