## Supplementary table 1 for "SOYBEAN RESPONSE TO INITIAL STAGE OF *Fusarium virguliforme* INFECTION"

**Supplementary table 1:** List of primers used for real-time quantitative RT-PCR (qPCR)

| **Name** | **NCBI identifier** | **Orientation** | **Sequence** | **Reference** |
| --- | --- | --- | --- | --- |
| PAL 1 | X52953 | Forward | CCAAGGAACCCCTATTGGAGCT | (Fernández et al. 2014) |
|  |  | Reverse | GTGTTGCTCAGCACTTTGGA |  |
| HCT | XM_003530175.1 | Forward | GGTCGCTCTCACACCCAATC | Mertz-Henning et al. 2015 |
|  |  | Reverse | CTCTCTGCTCGACTGCTCAC |  |
| CAD | AK286580.1 | Forward | ACGCAACATCCATCCTTCCA | Mertz-Henning et al. 2015 |
|  |  | Reverse | AACGTCGTGTAACCAAAACAGA |  |
| SKIP16 | CD397253.1 | Forward | GAGCCCAAGACATTGCGAGAG | Hu et al. 2009 |
|  |  | Reverse | CGGAAGCGGAAGAACTGAACC |  |

Fernández, I., Merlos, M., López-Ráez, J., Martínez-Medina, A., Ferrol, N., Azcón, C., et al. (2014). Defense Related Phytohormones Regulation in Arbuscular Mycorrhizal Symbioses Depends on the Partner Genotypes. *Journal of chemical ecology*, *40*, 791–803. doi:10.1007/s10886-014-0473-6

Hu, R., Fan, C., Li, H., Zhang, Q., & Fu, Y. F. (2009). Evaluation of putative reference genes for gene expression normalization in soybean by quantitative real-time RT-PCR. *BMC Molecular Biology*, *10*(May 2014), 93. doi:10.1186/1471-2199-10-93

Mertz-Henning, L. M., Nagashima, A. I., Krzyzanowski, F. C., Binneck, E., & Henning, F. A. (2015). Relative quantification of gene expression levels associated with lignin biosynthesis in soybean seed coat. *Seed Science and Technology*, *43*(3), 445–455. doi:10.15258/sst.2015.43.3.13
